# Genome-wide detection of genes under positive selection in worldwide populations of the barley scald pathogen

**DOI:** 10.1101/223545

**Authors:** Norfarhan Mohd-Assaad, Bruce A. McDonald, Daniel Croll

## Abstract

The coevolution between hosts and pathogens generates strong selection pressures to maintain resistance and infectivity, respectively. Genomes of plant pathogens often encode major effect loci for the ability to successfully infect a specific host. Hence, heterogeneity in the host genotypes and abiotic factors leads to locally adapted pathogen populations. However, the genetic basis of local adaptation is poorly understood. We analyzed global field populations of *Rhynchosporium commune*, the pathogen causing barley scald disease, to identify candidate genes for local adaptation. Whole genome sequencing data generated for 125 isolates showed that the pathogen is subdivided into three genetic clusters associated with distinct geographic and climatic regions. Using haplotype-based selection scans applied independently to each genetic cluster, we found strong evidence for selective sweeps throughout the genome. Comparisons of loci under selection among clusters revealed little overlap, suggesting that ecological differences associated with each cluster led to variable selection regimes. The strongest signals of selection were found predominantly in the two clusters composed of isolates from Central Europe and Ethiopia. The strongest selective sweep regions encoded proteins with functions related to both biotic and abiotic stresses. We found that selective sweep regions were enriched in genes encoding functions in cellular localization, protein transport activity, and DNA damage responses. In contrast to the prevailing view that a small number of gene-for-gene interactions govern plant pathogen evolution, our analyses suggest that the evolutionary trajectory is largely determined by spatially heterogeneous biotic and abiotic selection pressures.

## Introduction

Pathogens pose a major threat to human health and global food security. Hosts and pathogens are often locked into strong antagonistic coevolution, which is a major force driving biodiversity. During coevolution, adaptation in either of the partners generates selection pressure on the other partner, which in turn evolves counter-adaptations (Brockhurst & Koskella 2013). Pathogens are considered to be at an advantage in the coevolutionary race as they often exhibit shorter generation times, higher mutation rates, and larger effective population sizes relative to their host (Kaltz & Shykoff 1998; Gandon et al. 2002; Hamilton et al. 1990). Under reduced levels of gene flow, this asymmetry can lead to locally adapted pathogen populations, i.e. situations where a pathogen population is better adapted to its sympatric host population than allopatric host populations (‘local’ vs ‘foreign’ host) (Kaltz & Shykoff 1998). In a heterogeneous environment, a diverse set of host populations connected by variable levels of gene flow will generate complex selection pressures on pathogen populations over space and time and strong frequency-dependent selection can lead to local extinction and recolonization events (Kaltz & Shykoff 1998). Hence, the degree of local adaptation exhibited by hosts and their pathogens will be shaped by multiple factors, including the supply of genetic variation and their respective population sizes (Gandon et al. 1996).

Studies of local adaptation in pathogens have been sparse. Laine (2005) observed a geographical mosaic in the degrees of local adaptation in *Plantago lanceolata* - *Podosphaera plantaginis* metapopulations, where the major factor shaping the extent of local adaptation was likely the rate of pathogen dispersal among host populations. A similar observation was reported earlier by Thrall *et al*(2002) in a study between the host *Linum marginale* and the fungal pathogen *Melampsora lini*. In the *Colletotricum lindemuthianum* - *Phaseolus vulgaris* pathosystem, the pathogen was shown to have weaker levels of population differentiation than its host, which likely contributed to local adaptation of pathogen populations (Sicard et al. 1997; 2007; Cattan-Toupance et al. 1998). Local adaptation is not only shaped by rates of gene flow, but also by heterogeneity in the abiotic environment such as differences in rainfall and temperature. Gorter *et al* (2016) coevolved the bacterium *Pseudomonas fluorescens* and its viral parasite bacteriophage Φ2 at three different temperatures and found that the bacteria and phages were more resistant and infectious at the temperature where they previously coevolved. Similarly, the coevolution of *Plantago lanceolata* - *Podosphaera plantaginis* was found to be tightly linked to climatic factors (Laine 2008). However, what environmental factors are conducive to local adaptation and the relative effect of individual selective agents remain poorly understood.

The process of local adaptation in agricultural ecosystem may differ from the process that operates in natural host-pathogens systems (Croll & McDonald 2016). Agricultural ecosystems usually exhibit homogeneous environments with genetically uniform hosts planted over large areas. Though planting a single elite crop variety is thought to improve crop yields, the massive monoculture in turn imposes strong directional selection for pathogens to specialize on a specific host genotype (Stukenbrock & McDonald 2008). Highly specialized agricultural pathogens cause significant damage on host plants and lead to large-scale losses. The Irish famine in the 19th century was caused by the attack of an oomycete pathogen specialized on potato clones that destroyed the food supply of one-third of the population (Scholthof 2007). Widespread planting of a single banana clone “Gros Michel” in Panama and Costa Rica led to the emergence of the specialized pathogen *Fusarium oxysporum* f. sp. *cubense* (Ploetz 2000). More recently, a new race of the wheat stem rust pathogen *Puccinia graminis* f. sp. *tritici* surmounted the widely deployed stem rust resistance gene *Sr31* (Pretorius et al. 2000).

With the exception of a few highly clonal crops such as bananas, most agroecosystems exhibit both complex environmental differences and diversified host genotypes at the scale of countries and continents (Bianchi et al. 2006). The resulting mosaic in host genotypes and ecological niches creates the opportunity for divergent selection among pathogen populations. Zhan and McDonald (2011) found evidence for local thermal adaptation across global populations of the wheat pathogen *Zymoseptoria tritici*. Temperature-dependent local adaptation was also observed among isolates of *Puccinia striiformis* f.sp. *tritici* (PST) collected in northern and southern France (Mboup et al. 2012). The northern PST population harbored all virulence genes necessary to infect wheat varieties deployed in the south, but it failed to invade the southern region due to a lack of adaptation to the warmer Mediterranean climate. In addition to climatic limitations, extensive use of fungicides can impose strong directional selection on pathogen populations and lead to local adaptation. Selection for fungicide tolerance likely led to locally adapted *Z. tritici* populations (Zhan et al. 2006). Although evidence for divergent selection can be found through careful assessment of phenotypes, direct identification of the genes underlying the process of local adaptation remains rare.

The genetic basis of pathogen local adaptation in agricultural ecosystems is generally not well understood, though many microbial pathogens should be highly tractable models for genomic analyses (Croll & McDonald 2016). Until recently, the identification of adaptive loci in plant pathogens was largely limited to population analyses of pathogenicity genes (Schürch et al. 2004; Stukenbrock & McDonald 2007). However, pathogens likely experience a multitude of biotic and abiotic selection pressures. Hence, many loci across the pathogen genome are likely to harbor genetic variation under selection. The analyses of loci under recent selection have been revolutionized by genome sequencing at the population level. A number of statistical tests were specifically designed to retrieve signatures of adaptive loci in large-scale datasets (Voight et al. 2006; Lao et al. 2007; Bigham et al. 2010; Qian et al. 2013; Xuanyao Liu et al. 2017). Many tests of selection focused on one of the following signatures of a positively selected allele: an increase in linkage disequilibrium (LD), shifts in allele frequency spectra, or higher than expected levels of population differentiation (Vitti et al. 2013). Tools implementing these statistics have been widely used to detect signatures of recent selection in various organisms including humans, flies, the malaria pathogen, and livestock animals (Duffy et al. 2015; Zhao et al. 2015; Martin et al. 2016; Garud et al. 2015). One of the widely-used approaches is to calculate integrated haplotype homozygosity (iHS), which estimates the decay of the extended haplotype homozygosity (EHH) between an ancestral and derived allele at each SNP position (Sabeti et al. 2002; Voight et al. 2006). The rationale for this method is that the increase in frequency of a beneficial mutation will occur too quickly for recombination to homogenize its genetic background. Hence, a positively selected allele will be embedded in a less common and longer stretch of homogenous chromosomal sequence compared to neutral loci residing in more common and shorter haplotypes. Analyses of EHH are particularly suited to detection of soft (*i.e*. incomplete) selective sweeps (Voight et al. 2006). An extension to within-population EHH analyses is the cross-population extended haplotype homozygosity (XP-EHH), which is a more powerful method to detect nearly fixed selective sweeps because it compares haplotypes in a pair of populations (Sabeti et al. 2002). A combination of iHS and XP-EHH analyses should lead to a comprehensive set of candidate loci underlying recent local adaptation.

*Rhynchosporium commune* is an important fungal pathogen that causes barley scald disease globally with particularly high prevalence in temperate regions with cool and moist winters (Linde et al. 2003; Brunner et al. 2007; Aoki et al. 2011). *R. commune* is a member of the host-specialized *Rhynchosporium* species complex that split from a common ancestor 1200-3600 years ago (Zaffarano et al. 2008; 2011). The center of origin of *R. commune* is likely in Scandinavia, because this region is the diversity hotspot of the pathogen, unlike many other pathogens of crops that were domesticated in the Fertile Crescent (Zaffarano et al. 2006; Brunner et al. 2007). Individual field populations of this sexual pathogen are highly diverse with an average of 76% of the global genetic diversity found within a single barley field (Linde et al. 2003; 2009). The genetic basis of adaptation to its host is poorly understood with a few exceptions. Three genes encoding necrosis inducing peptides (*NIP1-3*)contribute to the development of necrosis (localized cell death promoting the infection) (Rohe et al. 1995; Wevelsiep et al. 1991; 1993; Kirsten et al. 2012; Stefansson, Willi, et al. 2014; Schürch et al. 2004) and a previously unknown pathogen-associated molecular pattern (PAMP) named Cell Death Inducing 1 (RcCDI1) is highly expressed during early infection of the host (Franco Orozco et al. 2017). *R. commune* populations can rapidly adapt to overcome host resistance (Xi et al. 2000) and were shown to adapt to abiotic selection pressures including temperature variation and fungicide applications (Stefansson et al. 2013; Brunner et al. 2016; Mohd-Assaad et al. 2016). A genome-wide association study (GWAS) revealed that resistance to azole fungicides emerged from mutations in multiple loci including the gene encoding the protein targeted by azoles (Mohd-Assaad et al. 2016).

A genome-wide analysis of nine global populations of *R. commune* showed that the species is subdivided into three main genetic groups (Mohd-Assaad et al. 2016). Each of these main groups was primarily associated with distinct geographic and climatic regions including Scandinavia, Central Europe and Oceania, and Ethiopia. Here we aim to identify candidate regions of divergent selection in these three genetic groups of *R. commune*. For this, we analyzed 125 whole-genome sequences collected from isolates across the distribution range of the pathogen. We performed genome-wide analyses of selection first at the population level using iHS and then at the between-population level using XP-EHH.

## Materials and methods

### Isolates collection, DNA preparation, and full genome sequencing

A total of 125 strains of *R. commune* were collected from nine countries: New Zealand, Australia, Ethiopia, Switzerland, Spain, Norway, Finland, Iceland, and USA (Figure 1A). Fourteen genetically distinct haplotypes were chosen from each field population with the exception of the USA population (n = 13). All isolates were previously characterized using microsatellite markers for population genetics studies (Stefansson, Willi, et al. 2014; Stefansson et al. 2013; Stefansson, McDonald, et al. 2014). We added isolates from two closely related *Rhynchosporium* species as outgroups, including nine isolates of *R. secalis* from Switzerland, France, and Russia as well as eight isolates of *R. agropyri* collected from different locations in Switzerland. Details for all *Rhynchosporium* spp. isolates used in this study are summarized in Supplementary Table 1.

**Figure 1:**
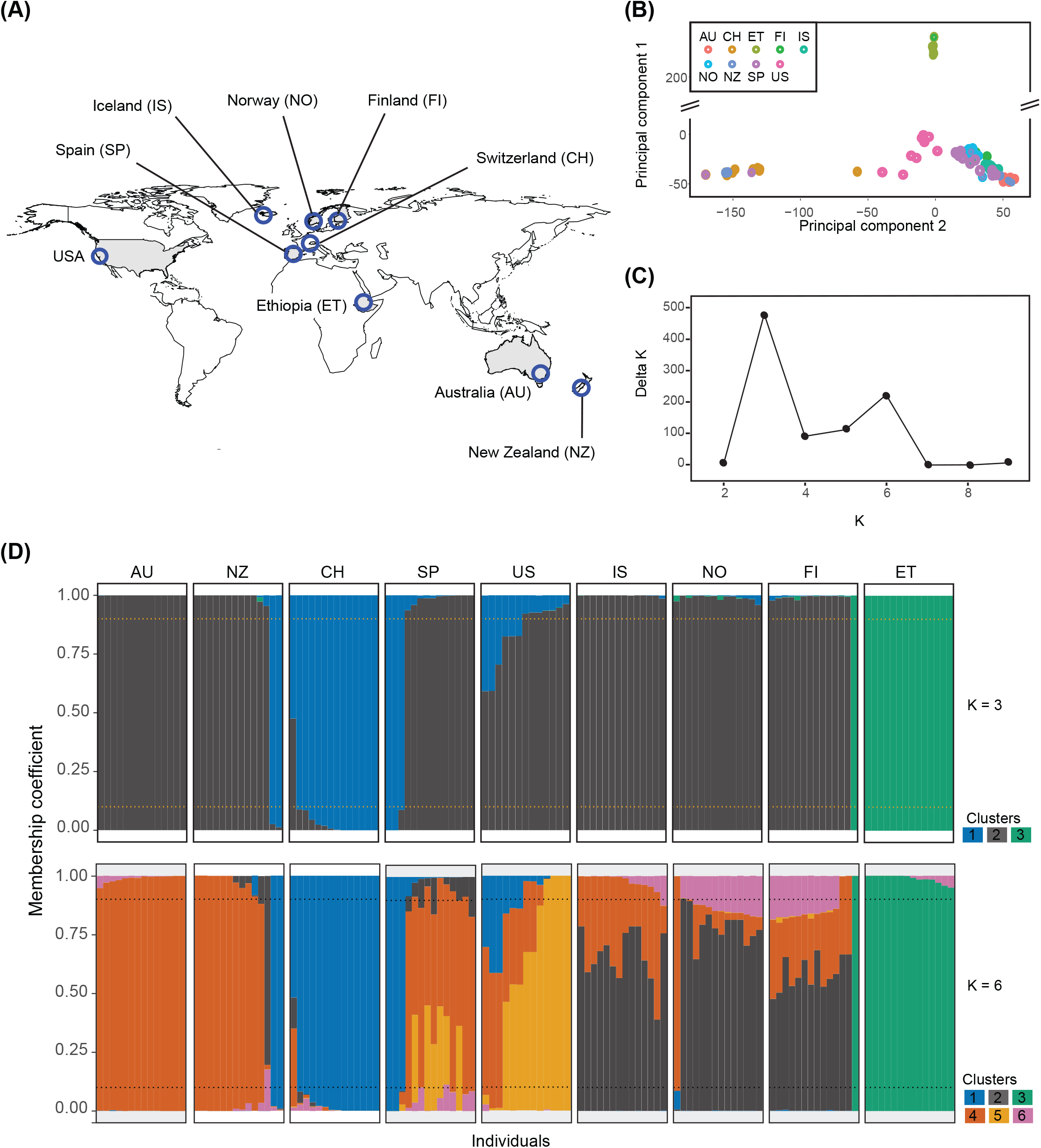
Population structure analyses of 125 *Rhynchosporium commune* isolates from a worldwide collection based on genome-wide single nucleotide polymorphisms. (A) Sampling locations of *R. commune* included in this study. (B) Principal component analysis of genetic differentiation among isolates. (C) Delta *K* plot of the STRUCTURE Bayesian clustering analyses for *Ks* of 2-9. (D) Genetic clustering using STRUCTURE for *K* = 3 (above) and *K* = 6 (below). Each colour represents one cluster and the vertical bar represents the proportion of cluster assignment for each genotype. Dotted lines are the minimum threshold value of membership coefficient for cluster assignment (90%).

Genomic DNA was isolated from mycelium grown in Potato Dextrose Broth (PDB) using DNeasy Plant Mini Kits (Qiagen) according to the manufacturer’s protocol. Paired-end sequencing of 125 bp reads was performed on a HiSeq2000 Illumina sequencer with an insert size of approximately 500 bp. The sequencing was performed at the Quantitative Genomics Facility (D-BSSE) of the ETH Zurich, Switzerland.

### Read mapping, SNP calling, and quality control

Low quality reads and traces of Illumina adapter sequences were trimmed using Trimmomatic version 0.32 (Bolger et al. 2014) with the following settings: trailing = 10, sliding-window = 4:10, and minlen = 50. Resulting high-quality reads were mapped to the reference genome of *R. commune* (Penselin et al. 2016) using Bowtie2 version 2.2.3 (Langmead et al. 2009). Aligned reads were flagged for duplicates using the MarkDuplicates program in Picard tools version 1.119 (http://broadinstitute.github.io/picard/).

Multi-sample single nucleotide polymorphism (SNP) calling was performed using two independent variant-calling software suites to ensure high accuracy: the Genome Analysis Toolkit (GATK) version 3.3-0 (McKenna et al. 2010) and Freebayes version v1.0.2-29-g41c1313 (Garrison & Marth 2012). For GATK, SNPs were called using the tools HaplotypeCaller and GenotypeGVCF. The maximum number of alternative alleles was set to two with a minimum phred-scaled quality score of 500. In addition, SNPs were selected if they matched the following criteria: 2 < BaseQualityRankSumTest < −2, 2 < ReadPosRankSumTest < −2, RMSMappingQuality < 30, 2 < MappingQualityRankSumTest < −2, QualByDepth < 20, and FisherStrand > 10. For Freebayes, we genotyped only SNPs and ignored insertion/deletions (indels), multi-nucleotide polymorphisms (MNPs) and complex events. Reads were required to have a minimum mapping quality of 30. The maximum number of alternative alleles was set to the best two alleles ranked by sum of supporting quality scores. SNPs with a phred-scaled quality score below 500 were removed from the dataset. SNPs called by GATK and Freebayes were compared using vcftools version 0.1.12b (Danecek et al. 2011). We only retained SNPs that were detected by both SNP caller suites for subsequent analyses. The joint SNP dataset was further filtered to retain only SNPs with a genotyping rate > 90%. Illumina reads generated from *R. secalis* and *R. agropyri* isolates were processed using the same procedure described above with the following exception: SNPs were retained if the genotyping rate was >50%. SNP genotypes retained for *R. secalis* and *R. agropyri* were used as outgroups to infer the ancestral state of SNPs in *R. commune*. A *R. commune* SNP allele was designated as ancestral if the allele was identified as fixed in both *R. secalis* and *R. agropyri*. Summaries of the read mapping, SNP calling and quality control pipeline are shown in Supplementary Figure 1.

### Prediction of gene functions

We predicted putative functions of all genes identified in the *R. commune* genome using InterProScan v.5.23-62.0 (Jones et al. 2014). The assignment of protein sequence motifs to protein families (Pfam) and gene ontology (GO) terms was performed based on hidden Markov models (HMM) implemented in InterProScan. Amino acid sequences were screened for evidence of secretion signals, and transmembrane, cytoplasmic and extracellular domains using a combination of SignalP v.4.1 (Petersen et al. 2011), Phobius v.1.01 (Käll et al. 2007) and TMHMM v.2.0 (Krogh et al. 2001). The *R. commune* secretome was defined as proteins predicted to include a secretion signal based on analyses using SignalP and Phobius. We removed any proteins with a predicted transmembrane domain based on Phobius and TMHMM analyses and a predicted cytoplasmic domain based on Phobius. We used the machine-learning approach implemented in EffectorP version 1.0 (Sperschneider et al. 2015) to identify the most likely effector proteins among the secreted proteins. We retained only predicted effectors with a posterior probability >0.8. All predicted secretomes were also screened for carbohydrate-active modules using the carbohydrate-active enzyme annotation (dbCAN) release 5.0 (Yin et al. 2012) for the identification of carbohydrate-active enzymes (CAZymes). We retained only protein hits with e-values <1e-17 and a coverage >45%. Repetitive elements in the reference genome of *R. commune* were annotated using RepeatModeler version 1.0.8 (Smit & Hubeley, RepeatModeler Open-1.0 2008-2015; http://www.repeatmasker.org). The repeat annotation was created by combining both matches to known repeat elements (Repbase version 20160629) and repeats identified *de novo* in the genome.

### Analyses of population structure

Genetic structure among *R. commune* samples was analysed using a principal component analysis (PCA) implemented in TASSEL version 20150625 (Bradbury et al. 2007). We also performed an unsupervised model-based Bayesian clustering implemented in STRUCTURE version 2.34 (Pritchard et al. 2000) to assign individuals to subgroups. To reduce the computational load, the SNP dataset used in STRUCTURE was reduced to one variant per 5 kb using the “--thin” option in vcftools, leaving 8,235 SNPs. The assignment of genotypes to clusters was run for total cluster numbers ranging from *K* = 1 to 10. We used the admixture model with correlated allele frequencies and no prior information about the demography. Each of the different *K*s was replicated 10 times with a burn-in period of 50,000 samples followed by 100,000 Monte Carlo Markov chain replicates. Parameter convergence was inspected visually. The output of the STRUCTURE analysis was processed using STRUCTURE HARVESTER (Earl & vonHoldt 2012) and the optimal number of subgroups was determined using the Δ*K* method (Evanno et al. 2005). Admixed isolates were identified as having a maximum membership coefficient <90%. These isolates were excluded from subsequent analyses of selection.

### Genome-wide scans for selection

Scans for selective sweeps were performed using two haplotype-based methods: integrated haplotype scores (iHS) and cross-population extended haplotype homozygosity (XP-EHH) implemented in the rehh package version 2.0.2 (Gautier et al. 2017) in R. For both analyses, we restricted our dataset to include only SNPs for which the ancestral allele was known (see above). The iHS analyses were performed independently for each of the genetic clusters identified with STRUCTURE using only SNPs that were polymorphic within the genetic cluster. Significant SNPs were defined by selecting the top 0.05% of |iHS| for cluster 1 and 3, and top 0.01% of |iHS| for the large cluster 2. For the XP-EHH analyses, we performed the test using all SNPs which were genotyped in at least 90% of isolates within each of the genetic clusters. SNPs were only retained if the ancestral allele was assigned (see above). The XP-EHH analyses were performed on pairwise comparisons of cluster 1 and cluster 3 against cluster 2 as the reference population unless stated otherwise. Significant SNPs were selected from the top 0.05% of |XP-EHH| for all clusters. We used default options for all analyses. However, we set the *maxgap* option to 20,000 whenever *calc_ehh*, *calc_ehhs*, or *scan_hh* was used. In addition to that, the threshold of missing data for haplotypes and SNPs was set to 90%.

Significant SNPs identified from either of the two selection scan methods and located on the same scaffold were grouped using a hierarchical clustering approach. If the distance between significant SNPs was below 50,000 bp, the SNPs were grouped into a single region under selection. Then, the extent of the region under selection was further refined by computing the extended haplotype homozygosity (EHH) for each significant SNP within a region under selection. The extent of the EHH above 0.05 was used to define windows around each SNP contained within a region and overlapping windows were then merged.

### Gene function enrichment analyses

Enrichment analysis of gene ontology categories was performed using the R package GSEABase and GOstats (Falcon & Gentleman 2007; Anders et al. 2015) with a false discovery rate set to 0.05. The minimum GO term size to be considered for enrichment analyses was set to at least five members in the reference genome. The enriched terms were then summarized by REVIGO (Supek et al. 2011) by removing redundant GO terms. We also analyzed evidence of enrichment in secreted proteins, effectors, cell wall degrading enzymes, and major facilitator superfamily (MFS) transporters using a hypergeometric test in R.

## Results

### Identification of high quality single SNPs with known ancestral state

We sequenced 125 *R. commune* isolates using Illumina to an average depth of 37X. We identified a total of 736,839 high quality SNPs using the GATK-HaplotypeCaller pipeline (Supplementary Figure 1). Then, we used FreeBayes to independently genotype all samples and found that 92% of the SNPs called by the GATK-HaplotypeCaller pipeline could be confirmed by FreeBayes (Supplementary Figure 1). We compared the performance of both methods based on several parameters. The SNP phred-scaled quality (QUAL) values and SNP allele frequencies called by GATK-HaplotypeCaller and FreeBayes were highly correlated (Supplementary Figure 2A, 2B). We also found a strong correlation of QUAL values and the alternative allele frequencies at the SNP loci. However, a small number of SNPs showed low QUAL values regardless of the alternative allele frequency (Supplementary Figure 2C). Interestingly, the physical positions of these SNPs were mainly in repeat-rich regions of the genome (Supplementary figure 2D). For all further analyses, we retained only SNPs identified by both SNP callers and genotyped in >90% of the isolates resulting in a dataset of 584,053 SNPs (Supplementary Figure 1). Isolates from the closely related sister species *R. secalis* (*n* = 9) and *R. agropyri* (*n* = 8) were sequenced to an average depth of 45X and 56X, respectively. SNPs sharing a fixed allele between the two sister species were used to assign the ancestral state of SNPs identified in *R. commune*. Ancestral states were assigned for 481,424 SNPs (82.4% of all retained *R. commune* SNPs). These SNPs were unevenly distributed along scaffolds (Figure 2B). Regions of high repeat density were largely devoid of callable SNPs due to our strict filtering procedure (Figure 2C).

**Figure 2:**
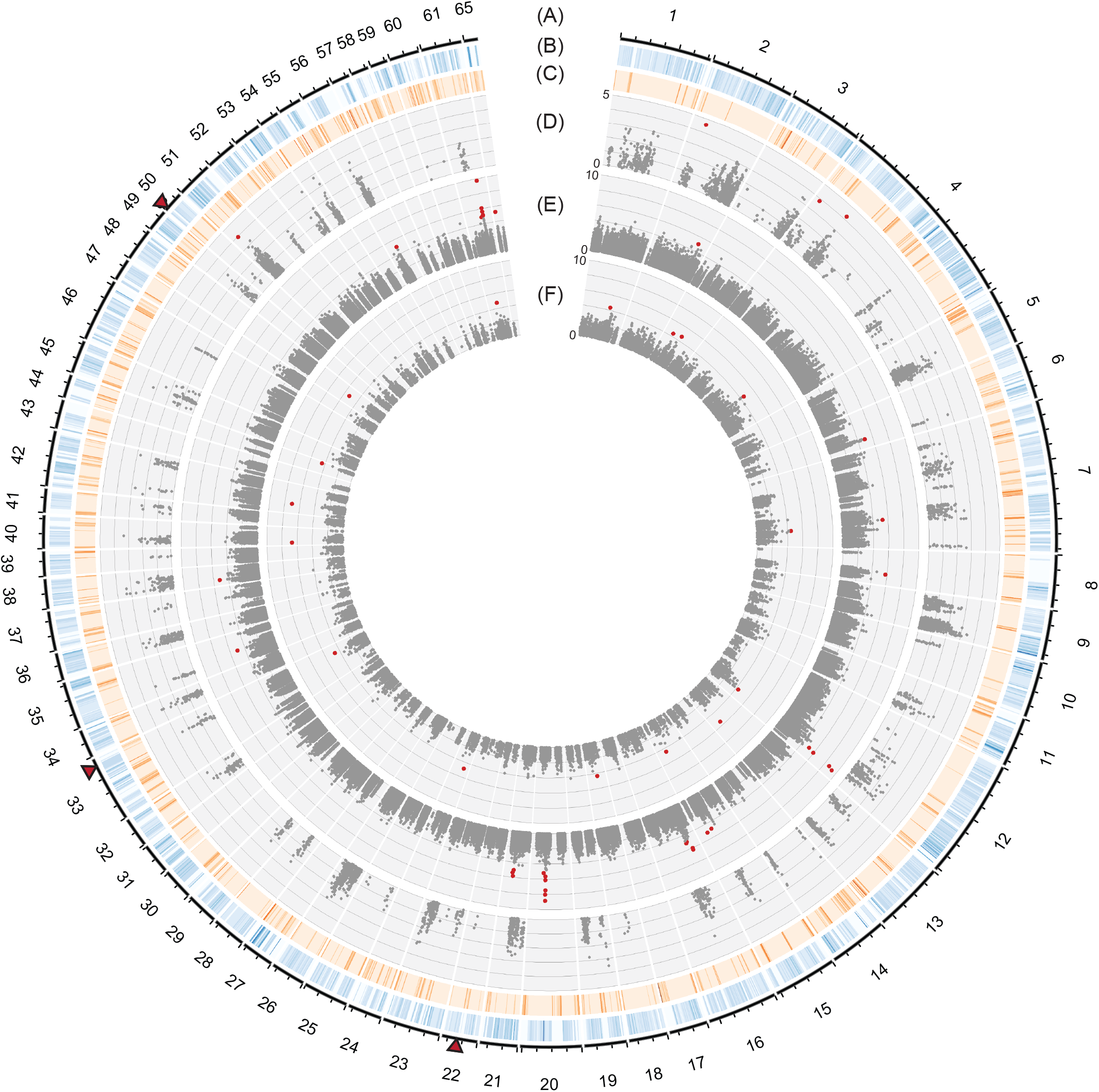
Genome-wide selection scans using the standardized integrated haplotype |iHS| score in three genetic clusters of *Rhynchosporium commune*. (A) Scaffolds of the reference genome and positions in Mb. (B) Density of single nucleotide polymorphisms (SNPs) across the whole genome shown in 10 kb non-overlapping windows (gradient shows coverage between 0-5%). (C) Coverage of repetitive elements across the whole genome shown in 10 kb non-overlapping windows (gradient shows coverage between 0-100%). (D-F) Dots show |iHS| values for each SNP polymorphic within the clusters 1, 2, and 3, respectively. The most significant SNPs (top 0.05% for clusters 1 and 3, and top 0.01% for cluster 2) were highlighted in red. Triangles near the outermost plot indicate the location of loci most significantly associated with resistance to the fungicide cyproconazole (Bonferroni threshold α = 0.05) in a genome-wide association study (Mohd-Assaad *et al* 2016).

### Population structure analyses based on genome-wide single nucleotide polymorphisms

We performed principal component (PC) and Bayesian clustering to analyse evidence for population subdivisions. Most populations including Scandinavia, New Zealand, Australia, and Spain were clustered into a single large group, whereas Ethiopian and Swiss populations formed two individual clusters from the main group (Figure 1B). The unsupervised Bayesian STRUCTURE analysis identified two probable genotype clustering scenarios at *K* = 3 and *K* = 6. *K* = 3 received the highest support consistent with the principal component analysis (Figure 1B, 1C). For *K* = 3, cluster 2 was found to be the most dominant cluster (Figure 1C, Figure 1D above). All genotypes from the Ethiopian population were assigned to a distinct cluster (Figure 1D). Genotypes from the Swiss population were mostly assigned to cluster 1, however several genotypes showed mixed ancestries from cluster 1 and 2 (Figure 1D above). Interestingly, two and three genotypes from New Zealand and Spain, respectively, were assigned to the cluster containing mainly Swiss isolates. In addition, a number of genotypes from the US population showed a mixed ancestry with contributions from cluster 1 and 2.

### Evidence for recent positive selection identified by integrated haplotype scores (iHS)

We screened for loci that experienced recent positive selection using EHH-based statistics. These statistics contrast the frequency of a haplotype to its relative EHH (defined from a core ancestral or derived allele). For all analyses, we used only SNP markers, for which the ancestral state could be assigned. First, we calculated the iHS statistics that compares the area under the EHH curve of the ancestral and derived alleles from the core allele. Performing the analyses separately for each genetic cluster, we identified a total of 39 genomic regions with signatures of selection across 27 scaffolds (Figure 2, Supplementary Table 2). The length of regions under selection ranged from 1,851 to 721,862 bp and contained 2 to 224 genes (Supplementary Table 2). We found that all regions under selection identified by iHS were each unique to an individual cluster. The highest percentage of identified loci were found in cluster 2 (46%), followed by cluster 3 (44%) and cluster 1 (10%).

To identify candidate genes under strong selection, we focused on the three most significant SNPs identified by iHS in clusters 2 and 3, whereas we focused on the two most significant SNPs in cluster 1, which generally showed lower levels of significance. We found a total of 12 candidate genes that were located within 5,000 bp of the most significant SNPs (Table 1). For cluster 1, we found two predicted candidate genes on scaffold 2, including the gene RCO7_04421 encoding a glucose-methanol-choline (GMC) oxidoreductase involved in plant cell wall degradation and RCO7_04423 encoding a multi-antimicrobial extrusion (MATE) multidrug transporter involved in the secretion of a wide range of metabolic and xenobiotic substances. In addition, we identified the gene RCO7_06862 on scaffold 4, encoding a KES1 oxysterol-binding protein with a role in the fungal ergosterol biosynthesis pathway. In cluster 2, we found the candidate gene RCO7_01309 on scaffold 61, which encodes an enzyme that catalyses the removal of an ammonia group from glutamine in a variety of substrates. The gene RCO7_02709 on scaffold 12 encodes a membrane-embedded UbiA prenyltransferase. In cluster 3, we found the candidate gene RC07_07301 on scaffold 42 encoding a concentrative nucleoside transporter (CNT), which mediates the uptake of nucleosides and nucleobases across the plasma membrane. We found no overlap between loci under selection and SNPs significantly associated with azole resistance in a previous GWAS analysis (Mohd-Assaad *et al* 2016).

### Positive selection identified by cross-population extended haplotype homozygosity (XP-EHH)

Unlike analyses of iHS comparing haplotype lengths within clusters, the XP-EHH test compares the haplotype lengths between pairs of clusters. We identified a total of 29 selective sweeps distributed across 19 scaffolds in all six pairwise cluster comparisons (Figure 3, Supplementary Table 2). Fifty-two percent of these selective sweeps were identified in cluster 3 while the remaining were shared evenly between cluster 1 and cluster 2. Interestingly, the selection targets on scaffold 1 (616,499-1,059,480 bp) and scaffold 9 (159,071-349,486 bp) were identified in two different pairwise comparisons (Figure 3, Supplementary table 2). In addition to that, a selection sweep on scaffold 2 (589,559-926,564 bp) was identified in cluster 1 and cluster 3 when XP-EHH was performed using cluster 2 as the reference cluster. Four of these selection sweeps (scaffold 1: 961,484-984,586 bp, scaffold 3: 623,832-625,683 bp, scaffold 12: 1,753,708-1,809,408 bp and scaffold 20: 533,694-710,548 bp) overlapped with selective sweeps identified using iHS. However, each of these selective sweeps were detected in different genetic clusters than the sweeps identified by iHS.

**Figure 3:**
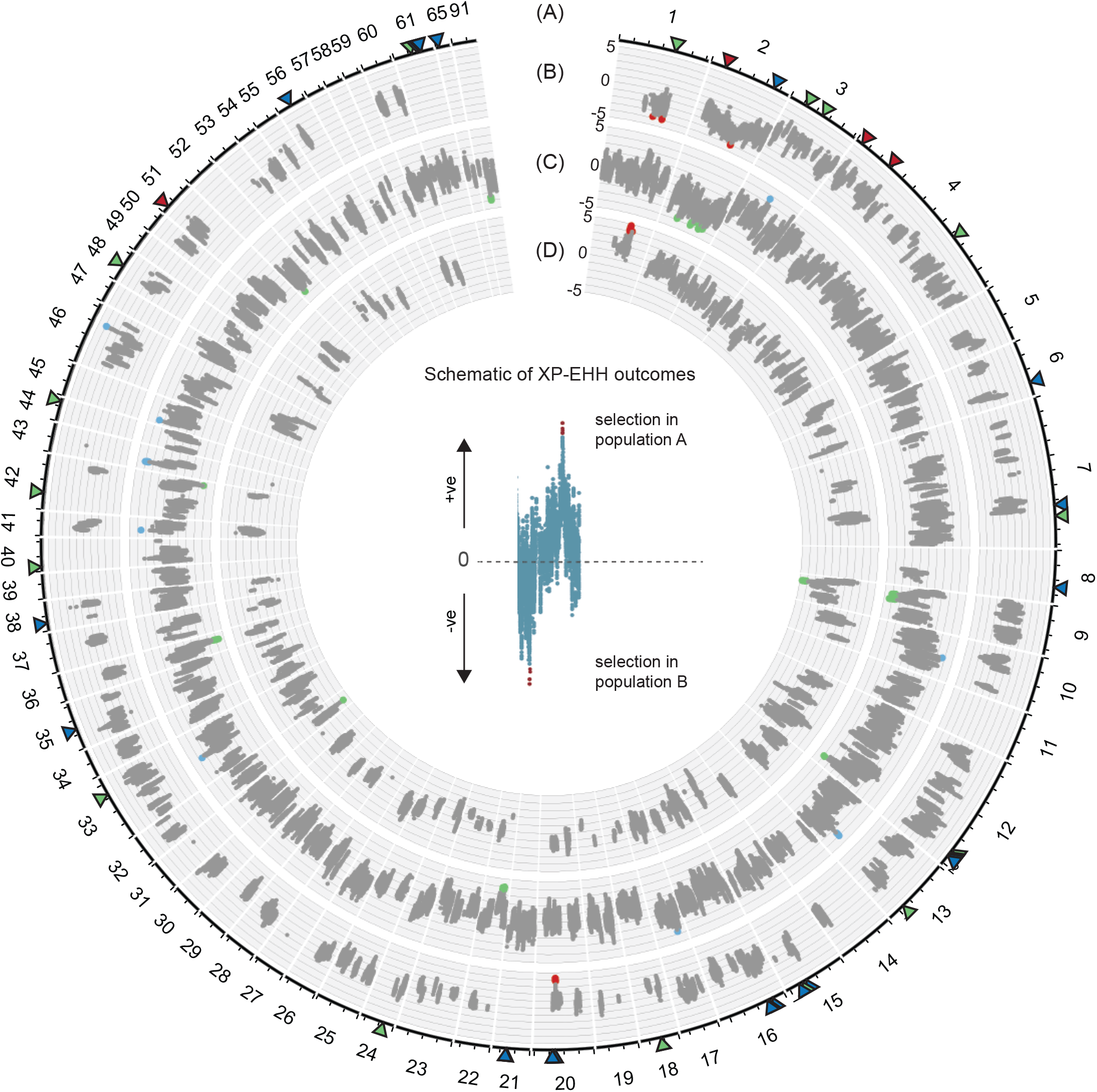
Genome-wide scan of the cross-population extended haplotype homozygosity (XP-EHH) analyses. (A) Scaffolds of the reference genome and positions in Mb. XP-EHH analyses of (B) cluster 2 vs cluster 1, (C) cluster 2 vs cluster 3, (D) cluster 1 vs cluster 3. The positive and negative values in panels B-D indicate the direction of selection. Positive values indicate positive selection in the reference population (cluster 2 was used as the reference population in panels B and C, cluster 1 in panel D). The most significant SNPs (top 0.05% of |XP-EHH|) are highlighted in red, blue, and green for signals of positive selection in cluster 1, 2, and 3, respectively. Triangles near the outermost plot show the most significant SNPs identified by the standardized integrated haplotype |iHS| analyses (red for cluster 1, blue for cluster 2 and green for cluster 3).

We focused on genes located in selective sweep regions identified both by XP-EHH and iHS analyses. We found a total of 10 genes that were located within 5,000 bp of the most significant SNPs identified by both XP-EHH and iHS analyses (Table 2). The two genes identified on scaffold 1 were RCO7_01116, encoding a phthalate 4,5-dioxygenase oxygenase reductase subunit involved in the degradation of xenobiotics, and an unknown protein encoded by RCO7_01117. We identified a ubiquitin-conjugating enzyme E2 encoded by RCO7_02480 and a MON2 protein encoded by RCO7_02479 on scaffold 3. MON2 plays an important role in endomembrane trafficking and Golgi homeostasis. We identified the RCO7_02696 gene on scaffold 12 that encodes a mediator complex subunit 7 (MED7) protein involved in the transcriptional regulation of nearly all RNA polymerase II-dependent genes. In addition to that, we identified RCO7_02697, which encodes a type 2 phosphatidic acid phosphatase (PAP2) gene important for lipid metabolism and signaling.

In order to identify additional candidate genes for local adaptation, we focused on selective sweep regions identified by two sets of cross-population tests. We found the gene RCO7_01132 located on scaffold 1 under positive selection in cluster 1 (using the cluster pairs 2-1 and 1-3 for XP-EHH analyses). This gene codes for a guanine nucleotide binding protein (G-protein) alpha chain, which plays an important role in various signaling systems. However, the two most significant SNPs identified on scaffold 9 in cluster 3 were not in proximity to any known gene. Using cluster 2 as the reference cluster, the strongest signal of a selective sweep was found in a large intergenic region. Using cluster 1 as the reference cluster, we identified RCO7_06601 encoding a Cu/Zn superoxide dismutase located 1,758 bp away from the top significant SNP. Cu/Zn superoxide dismutases play a role in protecting cells from damage caused by oxygen-mediated free radicals. Finally, we found two candidate genes on scaffold 2 shared by clusters 1 and 3 (when compared against cluster 2). The first candidate gene was RCO7_04583, encoding a transmembrane gene with a MARVEL domain which plays a role in membrane apposition events. The function of the second gene (RCO7_04584) is unknown.

### Gene functions overrepresented in selective sweep regions

We found a total of 972 and 811 genes in selective sweep regions detected by iHS and XP-EHH analyses, respectively. We analyzed whether genes under selection were enriched for specific functions by performing a gene ontology (GO) enrichment analysis. We found 16 terms that were significantly enriched (*p* < 0.01) for different functional categories (Figure 4, Supplementary Figure 3, Supplementary Table 3). Enriched GO terms did not overlap between gene sets identified by iHS and XP-EHH analyses, respectively. In iHS analyses, we found enrichment for localization (i.e. any process involved in the establishment and maintenance of cellular location) (GO:0051179), transport (GO:0006810), DNA damage checkpoint (GO:0000077), and DNA metabolic process (GO:0006259). Interestingly, we also found strong enrichment of the term reproduction (GO:0000003, *p* = 0.0001) and reproductive processes (GO:0022414, *p* = 0.0001) in regions under selection identified by XP-EHH. We investigated the genes encoding functions assigned to the GO term reproduction and related functions and found that these genes were related to meiosis and plasma membrane fusion, included a meiosis specific protein Spo22/ZIP4/TEX (RCO7_00959), a plasma membrane fusion protein PRM1 (RCO7_01086), and a DNA repair protein RAD51 (RCO7_01125). We found no evidence for enrichment in genes encoding secreted proteins, effectors, plant cell wall degrading enzymes or MFS transporters in regions under selection (Supplementary Table 4).

**Figure 4:**
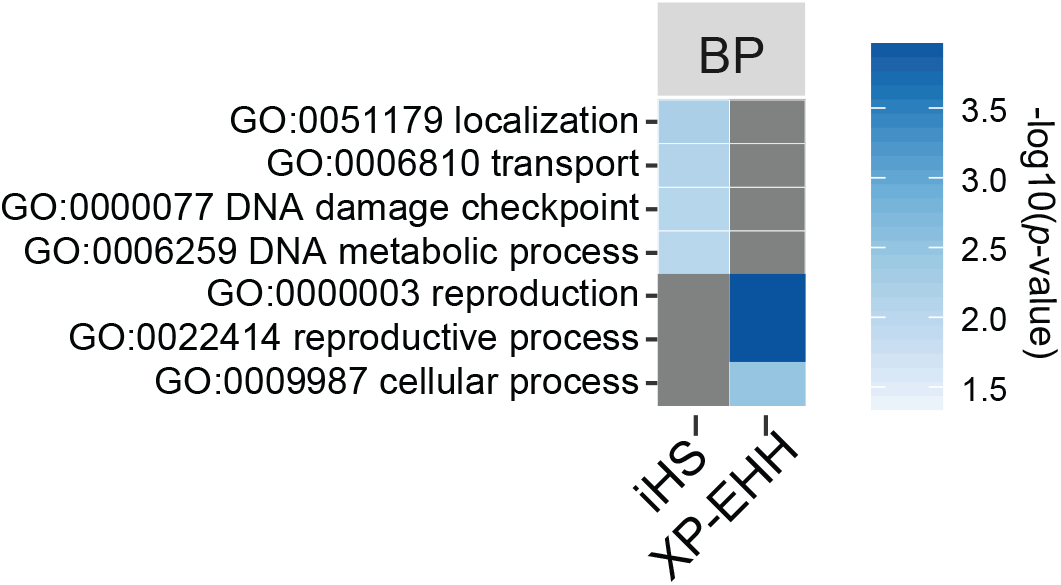
Gene ontology (GO) enrichment analyses for genes within selective sweep regions. Enriched GO terms related to biological process (BP) are shown for analyses of the integrated haplotype score (|iHS|) and cross-population extended haplotype homozygosity (XP-EHH). Loci were combined for all individual analyses of the three genetic clusters. GO terms were simplified using REVIGO prior to enrichment analyses and only significantly enriched terms *p* < 0.01 are shown.

## Discussion

We used population whole-genome sequencing to identify the signatures of selection in populations of the barley scald pathogen. Using two different haplotype-based selection scans, we found widespread signals of divergent selection in three genetic clusters of *R. commune*. The strongest signatures of selection were predominantly found in two of the three clusters. Contrary to expectations for a host specialized pathogen, loci showing strong evidence for selection were largely unrelated to adaptation to the host plant. Instead, we found enrichment in genes encoding functions in cellular localization, protein transport activity, DNA damage response and general metabolism. Genes related to reproductive processes were found to be significantly enriched in selective sweeps identified by cross-population analyses.

The global collection of the pathogen *R. commune* showed strong evidence for subdivision into three genetic clusters. Most of the genotypes were assigned to a predominantly northern European cluster. This cluster also comprised the genotypes collected near the center of origin of the pathogen (i.e. Scandinavia) (Zaffarano et al. 2006). Previous findings based on multilocus sequence data showed that *R. commune* likely expanded across the world from its center of origin (Zaffarano et al. 2009; Brunner et al. 2007). Nevertheless, contemporary gene flow was estimated to be low among continents. Our analyses of genome-wide SNPs confirmed the strong subdivision among continents. For example, the Swiss and Ethiopian populations showed striking genetic differentiation from the main cluster and each constituted largely independent clusters. However, multiple genotypes showed admixture between clusters suggesting ongoing genetic exchange. Interestingly, two and three genotypes of New Zealand and Spain, respectively, were assigned to the Swiss genetic cluster (>90% posterior probability), suggesting recent migration events and the absence of significant admixture. The US population showed frequent admixture from Swiss and Scandinavian populations showing that these two donor populations are playing an important role in the spread of *R. commune* to the North American continent. Previous studies suggested that gene flow from Europe was facilitated by the introduction of infected seeds during European colonization and may be ongoing to the present day (Brunner et al. 2007; Zaffarano et al. 2008; McDonald 2015). Despite evidence for migration events, gene flow was severely restricted among the genetic clusters. This should favor the emergence of local adaptation.

We performed genome-wide selection scans independently for each of the three genetic clusters and excluded admixed genotypes to avoid confounding effects of population structure. We found widespread evidence for selective sweeps across the genome, yet there was little overlap in loci under selection among clusters. The divergent selection pressures among clusters strongly suggests that differences in ecological factors imposed different selection regimes. For instance, in Ethiopia barley is grown over a broad climatic and edaphic range (barley is grown at altitudes from 1,400 to over 4,000 m above the sea level). Nevertheless, the pathogen is exposed to less seasonal temperature fluctuations than in Scandinavia (Asfaw 2000; Stefansson et al. 2013). Similarly, the climate in Oceania covers different seasonal fluctuations than Central European localities. Furthermore, our samples were isolated from genetically diverse barley landraces in Ethiopia (Hadado et al. 2010) while the Scandinavian and Swiss clusters comprised pathogen populations that were exposed to genetically homogeneous elite barley cultivars. Additional differences in environmental conditions include the early application of azole fungicides in Europe. Azole resistance was detected in many fungal pathogens in Europe, including *R. commune*, since the late 1990s (Cools et al. 2013). The Swiss population of *R. commune* already accumulated a number of azole resistance mutations as early as 1999 (Brunner et al. 2016; Mohd-Assaad et al. 2016). Taken together, the three genetic clusters comprised populations exposed to quite different environmental conditions providing substantial opportunity for local adaptation.

Our scans for selective sweeps in each of the three genetic clusters using iHS revealed a large number of loci under recent selection. These loci were broadly distributed across the genome. Importantly, we found little overlap in loci among the major genetic clusters, strongly suggesting differences in selection pressures among these clusters. Because *R. commune* has a well assembled and annotated genome, we investigated genes in immediate proximity to each of the selective sweep regions. We focused on the strongest signals of selection in each cluster and retrieved a total of 12 genes adjacent to these sweep regions. Half of these genes coded for proteins with a conserved function, including two membrane-bound transporters. The first MATE family transporter is involved in various metabolic pathways (Moriyama et al. 2008; Kuroda & Tsuchiya 2009; Juge Liu et al. 2016). In the rice pathogen *Pyricularia oryzae*, a MATE-family pump regulates glucose assimilation, sporulation, and pathogenicity (Fernandez et al. 2012). The second transporter mediates the uptake and release of nucleosides (Young et al. 2013). It is unclear what selection pressure would act on such a highly conserved transporter. Previous studies showed that nucleosides transporters can help attenuate nitrogen starvation, restore nucleotide pools and regenerate mycelium growth in nitrogen-free medium (Dean et al. 2014; Hamari et al. 2009; Daumann et al. 2016). Nucleoside and nitrogenous compounds salvation are important for pathogenicity in *Candida albicans, Xenorhabdus nemaptophila* and obligate parasitic protozoan (Orchard & Goodrich-Blair 2005; de Koning et al. 2005; Chitty & Fraser 2017). Additional genes in immediate proximity of selective sweeps included a gene encoding glucose-methanol-choline (GMC) oxidoreductase contributing to the degradation of lignin, a component of plant cell walls (Takahashi et al. 2015).

Fungicide resistance, in particular to azoles, evolved repeatedly in many plant pathogens (Cools et al. 2013). Several of the *R. commune* populations analyzed in this study were exposed to azoles and multiple loci were shown to confer increased resistance (Brunner et al. 2016; Mohd-Assaad et al. 2016). Azole resistance was found to be particularly high among isolates assigned to cluster 1. Interestingly, we identified a gene encoding the KES1 oxysterol-binding protein to be under selection in cluster 1. Mutations in the *KES1* gene can result in hypersensitivity to fungicides such as azoles that target ergosterol biosynthesis (Fang et al. 2012). Nevertheless, we found no evidence for positive selection at the loci previously associated with azole resistance by GWAS (Mohd-Assaad et al. 2016). This may be due to a combination of factors. First, fungicides were applied only for the last 30–40 years and the number of generations that has elapsed may have been too short for a strong selection signal to arise. Second, previous studies showed that azole resistance is a polygenic trait with possibly dozens of loci contributing to the overall level of resistance (Mohd-Assaad et al. 2016) making the detection of selective sweeps at individual loci less likely. Third, many mutations that conferred higher levels of resistance were also found in sensitive populations, suggesting that azole resistance at least partially arose from standing genetic variation. Fourth, some resistance mutations had negative pleiotropic effects on growth rates (Mohd-Assaad et al. 2016). Negative pleiotropy would weaken the response to selection due to fungicide exposure. Furthermore, the pathogen is only exposed to significant levels of fungicides during the crop growing season. Hence, resistance mutations with negative pleiotropic effects are likely under fluctuating selection which is not expected to generate strong signatures of selection.

In addition to our scans of recent positive selection within each genetic cluster, we used XP-EHH among pairs of clusters to gain sensitivity in the detection and to identify differences in signals among clusters. We focused in particular on loci that were confirmed to be under selection in a specific cluster by analyzing both pairwise comparisons with the other two clusters. We found such cross-validated loci for clusters 1 and 3. A gene encoding a guanine nucleotide binding protein was under strong selection specific to cluster 1. This protein is a part of heterotrimeric complex which has wide ranging roles in signaling, including known effects on reproduction, developmental processes and virulence in fungi (Gronover et al. 2001; Horwitz et al. 1999; Yamagishi et al. 2006; Tan et al. 2008; Gao & Nuss 1996). In cluster 3, a cross-validated gene encodes a Cu/Zn superoxide dismutase. These proteins enable pathogens to withstand reactive oxygen species (ROS), which are among the primary defense responses of plants during attack by pathogens (Yao et al. 2016; Rolke et al. 2004). Given their strong signatures of selection unique to individual clusters, both of these genes are strong candidates for genes affecting underlying processes of local adaptation for this pathogen. However, highly differentiated genomic regions can also arise in absence of local adapation (Bierne et al. 2011). Underappreciated environment-independent factors such as pre- and post-zygotic isolation or epistasis can lead to strong reductions in gene flow at specific loci. If such factors coincide with ecological variation, local adaptation is expected to produce indistinguishable signatures at loci under selection. However, we analyzed complete genomes instead of a relying on marker techniques that generate only reduced marker sets. Hence, we were able to analyze putative functions within selective sweep regions and found that many loci were indeed likely to encode functions related to biotic or abiotic adaptation. Analyzing the contribution of individual loci to phenotypic traits will enable a clear distinction between loci underlying local adaptation and endogeneous barriers.

Genes identified in selective sweep loci were only weakly connected with the repertoire of genes usually associated with highly specific host-pathogen interactions. Genes underlying host specialization often encode small secreted proteins. Such proteins directly or indirectly interact with a host receptor and modulate the host physiology to facilitate infection (Selin et al. 2016). We also found no evidence that genes encoding additional essential functions for plant pathogens such as cell wall degrading enzymes, proteases, lipases and MFS transporters were over-represented among selective sweep loci. These proteins play critical roles in the degradation of cell walls (Kikot et al. 2009; Esquerré-Tugayé et al. 2000), the secretion of virulence factors or detoxification (Coleman & Mylonakis 2009). In contrast, we found that selective sweep loci were overrepresented in genes encoding functions in transport, cellular localization and DNA damage repair. Such functions are more likely to be associated with selection pressures exerted by the abiotic environment (e.g. tolerance of external stresses such as UV). Genes in loci detected by XP-EHH were overrepresented in functions related to reproduction and meiosis. However, these genes were predominantly encoded on a single scaffold and hitchhiking selection may be an alternative explanation for this enrichment.

Similar to our study, Badouin *et al* (2017) found widespread selection across the genome of anther-smut fungi. Given the obligate association of anther-smut fungi with their hosts, selection pressure exerted by the host is likely to have a stronger impact on the pathogen. However, analyses of selective sweep loci revealed a multitude of possible targets not necessarily related to selection by the host. A number of studies focused on individual pathogenicity loci that encoded proteins that modulate the host immune system and may be detected by the host. Gene sequences encoding these proteins were found to show strong signatures of positive selection based on excessive non-synonymous vs. synonymous substitution rates (dN/dS ratio). (Schürch et al. 2004; Stukenbrock et al. 2011; Stukenbrock & McDonald 2007; Stukenbrock et al. 2010; Guyon et al. 2014). However, signatures of selection based on substitution rates arise over much longer time frames and reveal different selection regimes between species rather than selection pressure within species. Haplotype-based scans for selection across the genome generally detect much younger selection pressures (Vitti et al. 2013; Plissonneau et al. 2017).

The emergence of plant pathogens in agricultural ecosystems is a primary concern to ensure food security. The evolutionary processes through which plant pathogens rapidly overcome resistance in host plants or become resistant to fungicides provide illuminating case studies for the rapid evolution of complex traits. Our study showed that selection pressures operating across the distribution range of a globally distributed pathogen are likely to vary extensively. We found that the loci under recent positive selection encoded a multitude of functions related to abiotic and biotic stress factors. We found no evidence that selection imposed by the barley host played a dominant role in shaping recent selection pressures. This is surprising given the fact that many pathogen populations are known to encode major effect loci for aggressiveness (Presti et al. 2015; de Sain & Rep 2015). Instead, selection on *R. commune* led to selection on a large number of loci likely contributing to the rapid evolution of polygenic traits. Identifying the adaptive value of individual adaptive mutations and dissecting the genetic basis of complex traits will lead to major advances in understanding rapid evolutionary processes in populations with large standing genetic variation.

## Acknowledgements

We carried out part of the research at the Quantitative Genomics Facility of the D-BSSE and Genetic Diversity Center (GDC) of ETH Zurich. NM was supported by the Ministry of Higher Education Malaysia (MOHE) and Universiti Kebangsaan Malaysia (UKM) under the SLAI scheme. DC is supported by the Swiss National Science Foundation (grant 31003A_173265).

## Data accessibility

Raw sequencing data is available from the NCBI Short Read Archive under the BioProject accession number PRJNA327656.

## Author contributions

NMA and DC conceived and designed the study, NMA analyzed the data, NMA and DC wrote the manuscript. BAM provided funding, advice and edited the manuscript.

